# Visual stimulus-specific habituation of innate defensive behaviour in mice

**DOI:** 10.1101/2020.05.11.087825

**Authors:** Azadeh Tafreshiha, Sven A. van den Burg, Kato Smits, Laila A. Blömer, J. Alexander Heimel

**Affiliations:** Netherlands Institute for Neuroscience

**Keywords:** Innate behaviour, Stimulus-specific habituation, Freezing, Defensive behaviour

## Abstract

Innate defensive responses such as freezing or escape are essential for animal survival. Mice show defensive behaviour to stimuli sweeping overhead, like a bird cruising the sky. Here, we found that mice reduced their defensive freezing after sessions with a stimulus passing overhead repeatedly. This habituation is stimulus-specific, as mice freeze again to a novel shape. This allows us to investigate the invariances in the mouse visual system. The mice generalize over retinotopic location and over size and shape, but distinguish objects when they differ in both size and shape. Innate visual defensive responses are thus strongly influenced by previous experience as mice learn to ignore specific stimuli. This form of learning occurs at the level of a location-independent representation.

## Introduction

Innate visual responses are observed across the animal kingdom. Animals instinctively freeze or flee when they spot a threat (De Franceschi et al., 2016; Eilam, 2005; Temizer et al., 2015; Yilmaz & Meister, 2015), and orient towards more attractive targets (Ewert, 1987; Hoy et al., 2016). Even human new-borns instinctively respond to looming objects (Orioli et al., 2018). Though these innate behaviours might be hard-wired, they habituate by repeatedly being triggered (Randlett et al., 2019; Rankin et al., 2009; Thompson & Spencer, 1966). This habituation can be a reduction of the behavioural response, indiscriminately of the evoking stimulus, but can also be selective to the sensory modality (Vogel & Wagner, 2005) or even to the specific stimulus (Finkenstädt & Ewert, 1988; Gutfreund, 2012; Peeke & Veno, 1973; Schleidt et al., 2011). It is essential for animals to learn to ignore innocuous stimuli, but still respond to real threats. Mice habituate their defensive reaction to a repeatedly looming stimulus (Salay et al., 2018), but whether they show stimulus-selectivity in their defensive response to visual stimuli is not known. Here, we adapted the mouse behavioural paradigm of De Franceschi and colleagues and passed dark stimuli overhead, similar to birds of prey cruising the sky (De Franceschi & Solomon, 2018). We repeated the passing stimuli in several sessions to study if habituation of the defensive behaviour occurs. We investigated if the defensive behavioural response itself is habituated, or if the habituation would be specific to the stimulus. This also allowed us to determine if habituation is specific to the retinotopic position of the stimulus, such as the adaptation to a looming stimulus observed in the superficial layer of the superior colliculus of head-fixed mice (Lee et al., 2020), or if habituation is occurring further downstream in the visual pathway.

## Results

### Mice habituate to stimuli passing overhead

We placed the mice in an experimental box covered by a monitor and recorded from underneath (**Fig. 1A-B**). After a short familiarization period, a black disc traversed the screen in about 3 s (De Franceschi et al., 2016). Each session consisted of 20 such passes from right-to-left or vice versa with a minute between trials. Freezing was scored as the absence of any movement for at least 0.5 s that started while the stimulus was passing. had passed. During the first session, mice responded to the overhead moving disc by freezing in half of the trials (**Fig. 1C-D**). We only observed an escape running response in the presence of a shelter in the box. Here, we only describe experiments without a shelter. We repeated this paradigm with the same stimulus for five sessions with one or two days between sessions. Mice strongly habituated to the sweeping stimulus, and freezing decreased quickly over sessions (**Fig. 1D**). The percentage of trials in which the mouse froze dropped from 49 % in the first session to 16 % in the fifth session (paired t-test p = 0.006, 7 mice). The freezing response and its subsequent habituation was not specific to the disc stimulus. We repeated the same test, with naïve mice, using a black silhouette of a hawk (**Fig. 1E**). This gave very similar results (**Fig. 1F**). In the first session, mice froze on 46 % of the presentations (not different from the disc, t-test p = 0.78, 7 mice each group). This dropped to 19 % in the fifth session, much smaller than in session 1 (paired t-test p = 0.021), and not different from the fifth disc session (t-test p = 0.65). The habituation is therefore not due to the artificial nature of the disc stimulus.

**Figure 1.**
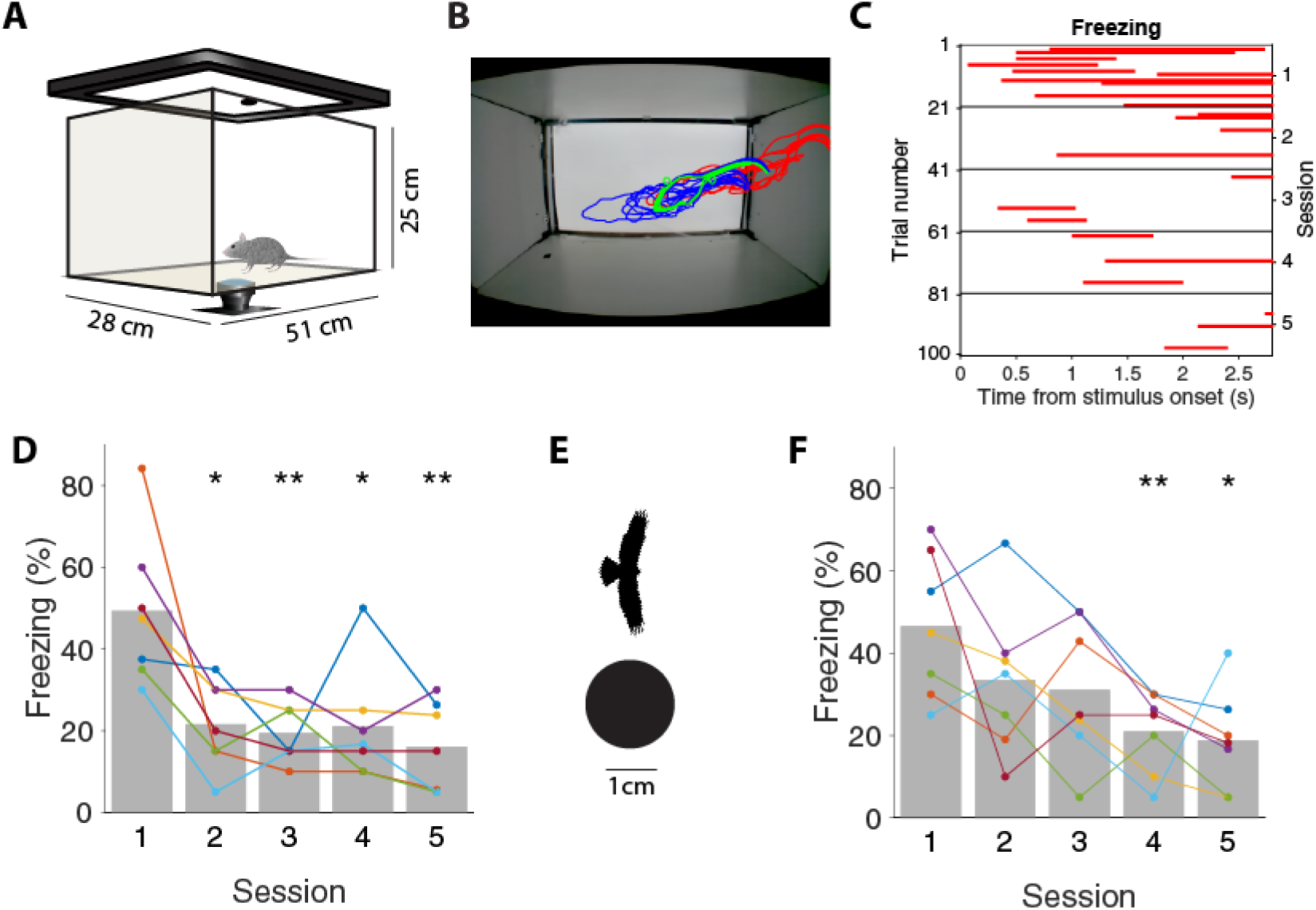
Mice habituate to stimuli passing overhead. A) Acrylic glass observation box with wide-angle camera mounted below and display mounted on top. B) Snapshots of mouse outline every 0.33 s. In red, the 3s before stimulus onset. In green during stimulus. In blue, 3s after the stimulus disappeared. C) Freezing periods for the 5 sessions of one mouse indicated in red. D) The percentage of freezing to a disc passing overhead for all trials in each session. Each colour represents a mouse. Paired t-tests of first session versus later sessions, 7 mice, *: p<0.05; **: p<0.01. E) The disc and hawk stimuli. F) Freezing for sessions with the hawk stimulus. Colours indicate mice (different mice from D). Paired t-tests of first session versus sessions, 7 mice, *: p<0.05; **: p<0.01.

### Habituation is not specific to stimulus location

The habituation in the combined group of mice took place gradually and the average amount of freezing decreased continuously over the five sessions. This gradual, but noisy, decrease was observed in the individual mice. The steady decrease of the mean amount of freezing was not caused by different mice showing sharp decreases in freezing in different sessions. One explanation for a gradual decrease in freezing could be that there is fast but stimulus location-specific habituation. The hypothesis would be that at a specific retinotopic position the stimulus would only lead to freezing once and then would be ignored when it would later appear at the location again, in the absence of any negative consequences at the first presentation. With repeated presentations, more and more of the retinotopic map would be covered and the amount of freezing would reduce to baseline levels when the whole visual field has been covered. To investigate this possibility, we determined the head direction of the mice by measuring the position of the snout relative to their centre of mass. Next, we computed the position of the stimulus relative to the mouse while it was moving across the screen above the mouse, and plotted when the mouse started freezing (**Fig. 2A**, freezing in red). Not surprisingly, we found that over sessions the stimulus had traversed most of the visual field (**Fig. 2B**). As the example in Figure 2B shows, the mouse was often frozen when the stimulus was behind its head. This was common to all mice. The reason for this is that mice spend much time investigating the walls and in particular the corners of the box, and were thus pointing away from the stimulus (**Fig. 2C**). However, the conditional probability of a freeze to start at a stimulus position relative to the head was, as expected, highest in front of the mouse, where the stimulus could be seen (**Fig. 2D**).

**Figure 2.**
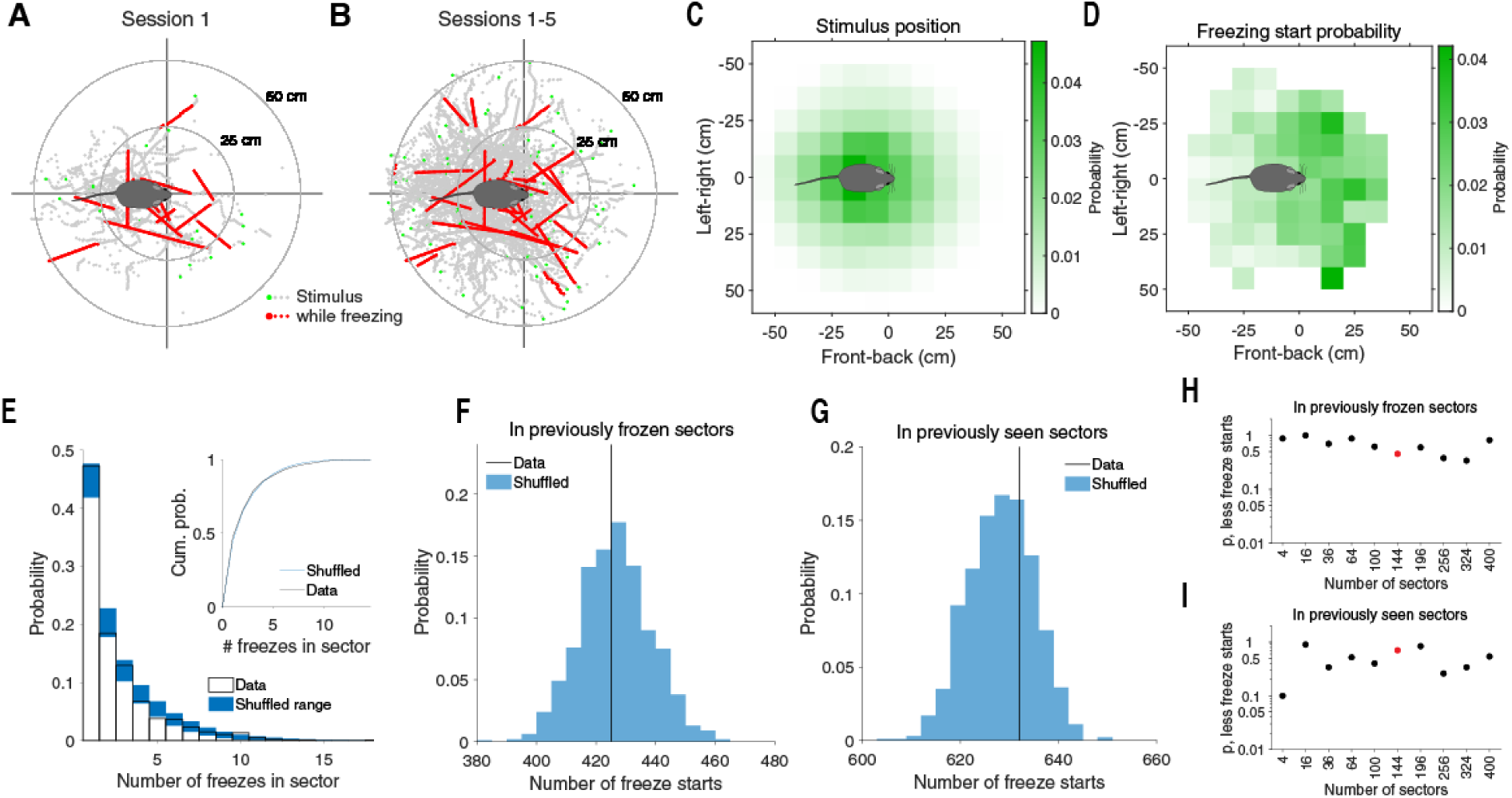
Habituation is not specific to stimulus location. A) Location of the stimulus relative to the mouse for the first session of an example mouse. The entry of the stimulus is shown in green. Red marks the stimulus trajectory while the mouse was freezing. Mouse is not drawn to scale. B) Same as A but for all 5 sessions. C) Probability density of the relative stimulus locations during the entire trajectories across all mice. Mouse is not drawn to scale. D) The conditional probability of a freeze to start for a given stimulus location. E) Distribution of the number of freezes occurring for each stimulus location for the division in sectors used in C and D. In blue the range of 2 standard deviations below and above the mean for the cases when the freezes are shuffled across mice. Inset shows the cumulative distribution of the data and mean of the shuffled distributions. F) The number of freeze starts that occur in a sector that was previously traversed by the stimulus while a mouse was frozen (black line) falls inside the distribution when the freezes are shuffled across mice. G) Same as F for sectors traversed by the stimulus regardless of whether the mouse was freezing. H) Fraction of number of freeze starts in previously frozen sector of shuffled distribution that is smaller than the observed number. Red marker indicates case shown in F. P-values are not corrected for multiple comparisons. I) Same as H for the number of freeze starts in previously seen sectors. Red marker indicates case shown in G. See also Figure S1.

If the habituation were stimulus location-specific, then mice would have been less likely to freeze again when the stimulus occurred at a location in the visual field to which the mouse previously froze. This would imply that the number of freezes per sector of stimulus locations over all sessions for one mouse would be less than if the freezes would occur randomly. We therefore split the overhead visual field in a number of stimulus sectors (as in **Fig. 2C-D**) and computed the distribution of the number of freezes per sector. This distribution was identical to the distribution that we obtained when we shuffled the stimulus trajectories of all trials in which a mouse froze across all mice (**Fig. 2E**). Furthermore, the number of freeze starts that occurred in head-relative location sectors that the stimulus previously crossed while the mouse was frozen was not different from the number one gets if freezes were shuffled across mice (**Fig. 2F**). Finally, the number of freeze starts in sectors that the stimulus previously traversed, regardless of whether the mouse was freezing or not, was not different from the shuffled cases (**Fig. 2G**). All this strongly suggests an absence of location-specific habituation. The exact number of freeze starts in **Figures 2E-G** depends on the size and shape of the sectors. When taking the sectors too large, then a finer location-specific habituation could be missed. When taking the sectors too small, then our inability to measure the exact gaze direction could remove evidence of location-specific habituation. However, changing the number of sectors (**Fig. 2H-I**) or changing to a spherical division of visual space (**Fig. S1**) does not change the conclusion that we found no evidence of stimulus location-specific habituation.

### Habituation is stimulus-specific

The absence of location-specific habituation suggested that the mice habituated to the experience of harmless stimuli passing overhead, regardless of having seen the stimulus before from a specific angle, or perhaps even regardless of having seen the specific stimulus before at all. To check if mice generalized habituation to all stimuli, we showed in the sixth session ten times the hawk and ten times the disc stimulus, in random order. Mice showed increased freezing again in response to the stimulus that was novel to them (**Fig. 3A-B**). To the novel stimulus, they froze in 39 ± 7 % (mean ± s.e.m.) of the passes, while in the same session they only froze to 18 ± 5 % of the passes of the habituated stimulus (paired t-test p = 0.0081, 14 mice). The freezing to the newly introduced stimulus was not significantly lower than the freezing in the very first session (48 ± 5 %, paired t-test p = 0.2). The behaviour of mice thus clearly indicated a difference between the habituated stimulus and the novel stimulus. There was no difference between the cases where the hawk or the disc was the habituating or novel stimulus (two-way ANOVA p = 0.42, 7 mice each group).

**Figure 3.**
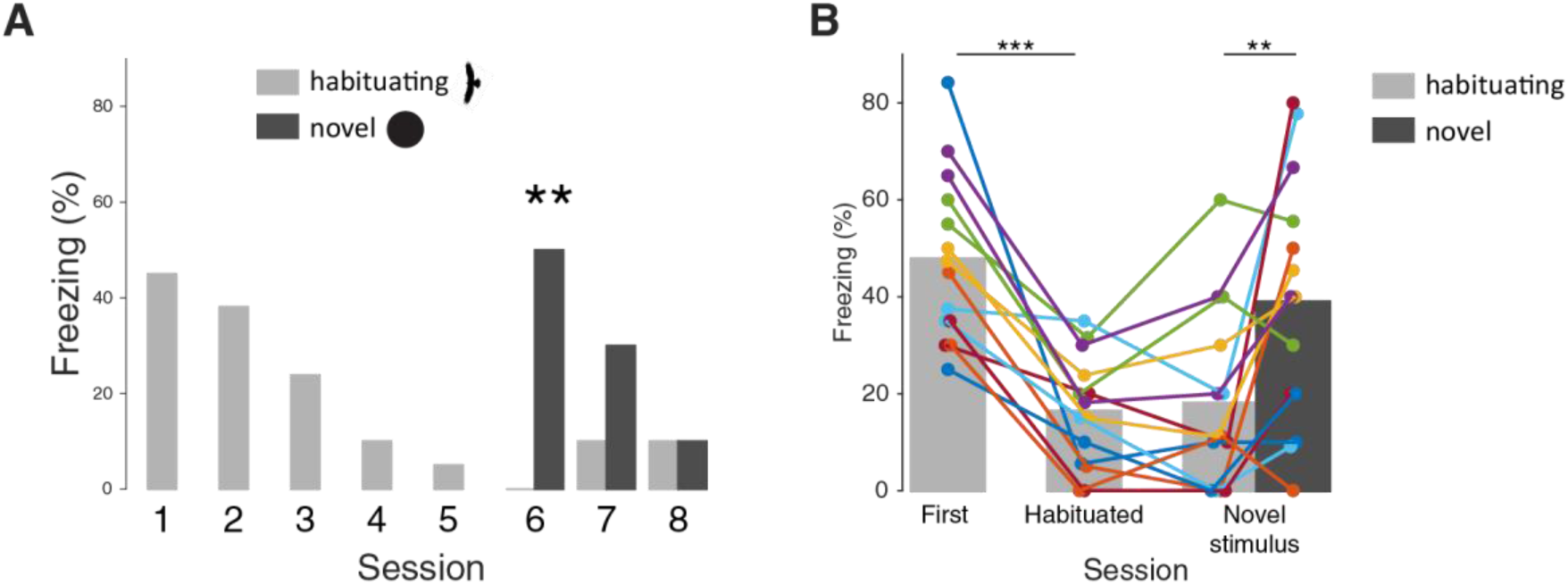
Habituation is stimulus-specific. A) Example of a mouse freezing in response to a habituating hawk stimulus and a later introduced novel disc stimulus (differences: session 6, chi^2^-test p = 0.0098; session 7, chi^2^-test p = 0.26; session 8, chi^2^-test p = 1; 10 trials for each stimulus in each session). B) Freezing in first session and last habituated session with a single stimulus, and the freezing in the first session when a novel second stimulus is introduced. Lines indicate mice. Difference habituating stimulus in first versus last session with single stimulus (paired t-test p = 4.2 × 10^−5^, 14 mice, ***). Difference novel versus habituating stimulus in first test (paired t-test p = 0.29). Difference novel versus habituating stimulus in same session (paired t-test p = 0.0081, **).

### Habituation is specific to combination of surface area and shape

We were interested in understanding which specific features alert a mouse that a novel stimulus has appeared. Taking into account that a hawk is a natural predator of the mouse, one could hypothesize that elements common to birds like feathers, tail and a beak distinguish the disc from the bird. However, given the limited acuity of mice (0.5 cycles per degree) (Prusky et al., 2000), the question was how much of these details they could actually perceive. Indeed, if we simply blur the stimuli to a degree that a grating at the acuity limit (0.5 cycles per degree) disappear, the bird decidedly starts looking like an ellipse (**Fig. 4A**). We therefore hypothesized that the mice would not be able to distinguish between the hawk and an ellipse that matched the hawk in surface area and length along its long axis. Naïve mice responded to the ellipse by freezing and again habituated to it over several sessions. However, when we then introduced the original hawk stimulus they would not freeze significantly more to the novel hawk stimulus than they would to the habituating ellipse stimuli. The same was true when we reversed this stimulus order in naïve mice. From the first session to the last session with a single stimulus, freezing was strongly reduced (paired t-test p = 0.00019, 9 mice, **Fig. 4B**), but in the session when the second stimulus was introduced, mice did not freeze more to the novel stimulus than to the habituated stimulus (p = 0.076). Therefore, in this freezing assay, mice did not distinguish the hawk and ellipse as different stimuli, indicating that avian features are not enough to make two stimuli distinguishable as overhead threats. Apparently, the hawk and the disc were distinguished by their other differences. They also differed in their aspect ratio and by a factor 2.5 in surface area. To separate these two features, we created two new stimuli (**Fig. 4C**): a smaller disc with the same surface area as the hawk, and a larger ellipse with the same aspect ratio as the hawk and the same surface area as the original disc.

**Figure 4.**
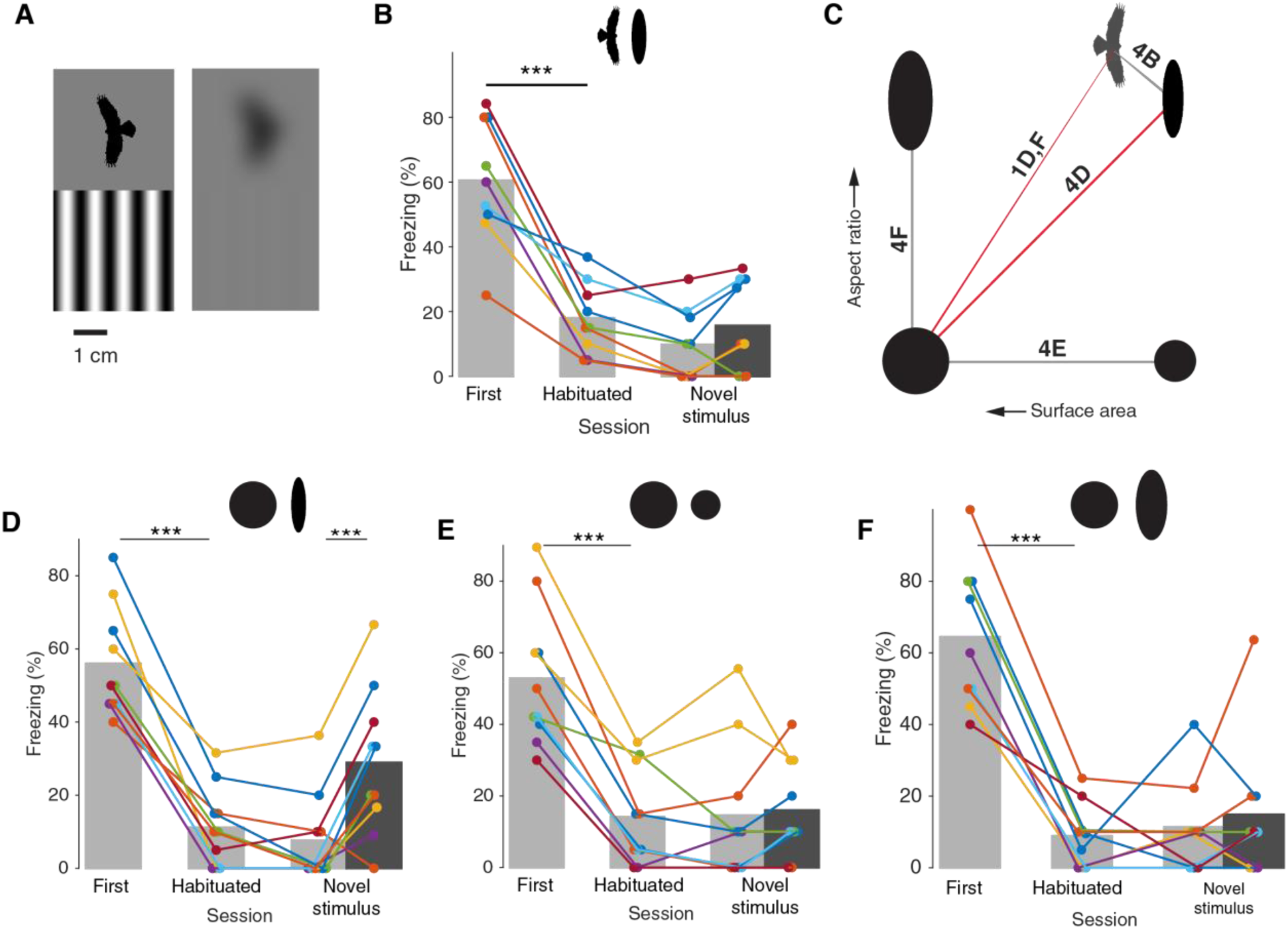
Habituation is specific to combination of surface area and shape. A) Left. Hawk stimulus and a sinusoidal grating at the mouse visual acuity limit (0.5 cycles per degree). Right. Same but after Gaussian blurring that annulled the sinusoidal grating. B) Freezing in sessions where either a hawk stimulus or an ellipse with an equal surface area and height were used. The animals habituated to the initial stimulus in five sessions (paired t-test p = 0.00019, 9 mice). No difference between the novel and habituated stimulus (p = 0.076). C) Stimuli were ellipses differing in surface area or aspect ratio. Hawk is shown next to closest matching ellipse. The large disc and hawk were used for the earlier figures. D) Mice habituate to a large disc or hawk-sized ellipse (paired t-test p = 4.9 × 10^−6^, 10 mice), but freeze again to the introduction of the other stimulus (p = 0.00081). E) Mice habituate to a large or small disc (paired t-test p = 1.8 × 10^−5^, 10 mice), but do not freeze again to the novel stimulus (p = 0.73). F) Mice habituate to a large disc and an ellipse with the same area (paired t-test p = 1.7 × 10^−5^, 9 mice), but do not freeze again to the novel stimulus (p = 0.57).

Next, we wanted to reproduce the original experiment of **Figure 3** with the original disc and the small ellipse. We repeated the entire paradigm with naïve mice, using the disc and the ellipse, distinguishable only by surface area and aspect ratio (**Fig. 4D**). Mice again habituated to either stimulus (paired t-test p = 4.9 × 10^−6^, 10 mice). When the novel stimulus was introduced after the habituating sessions, the mice clearly increased freezing again in response to the novel stimulus compared to the habituated stimulus (p = 0.00081). Mice thus showed stimulus-specific habituation also in this case, demonstrating that they distinguished two shapes when they were different in surface area and aspect ratio.

Concluding that mice perceived a difference between these stimuli, we were then interested if any stimulus parameter alone caused the perception of this difference. To test for surface area, we habituated new naïve mice with either the original disc or the smaller disc. While mice habituated to both discs as in earlier experiments (first versus last single-stimulus session, paired t-test p = 1.8 × 10^−5^, 10 mice), there was no difference in freezing to the disc with the habituated size and the one with the novel size (p = 0.73, **Fig. 4E**).

Finally, we tested if the difference in aspect ratio alone was sufficient for the stimuli to be distinguished. For this experiment, we used an ellipse matching the original disc in surface area. We repeated the behavioural paradigm in naïve mice. Mice showed freezing and habituation to both stimuli (different between first and last session with one stimulus, paired t-test p = 1.7 × 10^−5^, 9 mice). However, they showed no sign of stimulus-specific habituation to these stimuli (novel vs habituating, p = 0.57, **Fig. 4F**). These results demonstrate that neither surface area nor aspect ratio alone were the distinguishing features for mice, but in the case of hawk versus disc, a combination of the two was.

## Discussion

Our results show that mice respond to visual threatening overhead stimuli, and that they adjust their behaviour as their experience with the environment accumulates. In particular, they stop freezing to specific stimuli after repeatedly being exposed to them.

Our experimental paradigm is similar to the 1937 hawk/goose experiment by Lorenz and Tinbergen (Lorenz, 1939; Schleidt et al., 2011; Tinbergen, 1939). They moved cardboard bird silhouettes over young birds to investigate how birds are able to distinguish between life-threatening raptors and harmless flying creatures. They famously confirmed the hypothesis by Oscar Heinroth that the turkey chicks distinguished raptors from birds like geese by their short neck. Although this was taken by Tinbergen as evidence for an innate template, the chicks had seen geese before and stimulus-selective habituation had probably occurred (Schleidt et al., 2011), exactly as we now find in laboratory mice.

We find that mice in their defensive responses generalize over the specific shape and size, but distinguish ellipses that vary in both aspect ratio and size. How the mouse visual system makes this distinction is unclear. It is highly unlikely that the distinction is made based on the pixelwise differences in the retinal image. The difference in the pixelwise distances between the large disc and small ellipse, which mice do distinguish, and the large disc and small disc, which they do not distinguish, is only around 5%. If the comparison was performed directly on the retinal images, one would expect much smaller differences in the freezing response to novel small ellipse and small disc after habituation to the large disc. Further evidence that the stimulus-specificity happens at a more abstract level than the physical image, is the lack of specificity to the head-relative location at which the stimulus appeared. Mice froze equally to stimuli at head-relative positions, which stimuli had traversed before, as well as at novel locations. We did not measure the rotation of the eye within the head, and therefore do not know the gaze-direction with a high accuracy. The stimulus paths shown for example in **Figure 2B**, however, show that the mice do not freeze to retinotopic positions that they had not seen before. Even if the stimulus trajectories were highly inaccurate, there were not enough frozen paths to cover the entire field of view and stimulus location-independent habituation must have been occurring.

In the rodent, these innate visual defensive responses are thought to be mediated by the superior colliculus (Dean et al., 1989). Its superficial layer receives direct input from the retina, and from the superior colliculus information travels directly and indirectly to the periaqueductal grey (Evans et al., 2018), where freezing is initiated (Tovote et al., 2016). The level of abstraction and lack of retinotopic specificity suggest that the habituating plasticity occurs down-stream of the strictly retinotopic superficial layers of the superior colliculus. In the stimulus-comparator theory of habituation (Sokolov, 1963), the brain builds a model of the stimulus, and the fit of the stimulus to the model inhibits the stimulus-response pathway. In the mouse, the model-building and comparison could be computed in the visual cortex, where stimulus-specific response potentiation occurs (Cooke et al., 2015). Alternatively, we might be witnessing a stimulus-unspecific habituation downstream in the pathway, in combination with a novelty detector, which enhances responses when a new stimulus is detected.

In the laboratory mouse, we can start to investigate the locus of this form of experience-dependent plasticity. The relevance of this mechanism is not confined to mice. Stimulus-selective habituation of defensive responses has been observed in different species and modalities (Deecke et al., 2002). Already in one of Aesop’s fables, written long ago, the fox stopped being frightened by a lion after seeing it two times (Thompson, 2009). In humans, innate defensive behaviours are less easily evoked, but do occur and habituate (Bradley et al., 1993). It is of interest to know how this happens under normal circumstances and in neuropsychiatric conditions, such as autism, ADHD, and post-traumatic stress disorder where habituation is abnormal (Lissek & van Meurs, 2015; McDiarmid et al., 2017).

## Materials and Methods

### Experimental animals

Male C57BL/6JRj mice from Janvier Labs (Saint-Berthevin, France) were used. We received the mice at p30. They were housed in pairs in a normal 12 hour day/night cycle with food and water available ad libitum. Experiments were performed in the day-part of the cycle. Animals were 5 to 8 weeks old at the first session. Experiments were performed in accordance with national guidelines and regulations, and approved by the institutional animal care and use committees of the Royal Academy of Arts and Sciences and the Netherlands Institute for Neuroscience.

### Experimental setup

The experimental setup consisted of a rectangular acrylic box (51 cm long, 29 cm wide, 25 cm high) on a raised platform (18 cm) with an open top, a transparent bottom and opaque white walls. An LCD display (Dell) matching the dimensions of the box was placed on the top of the box to show the stimuli. We filmed the experimental session with a Raspberry Pi Camera outfitted with an 0.4x wide-angle lens placed underneath the setup. After each experiment, the inside surfaces were cleaned with 50% Ethanol and then water to remove any leavings and odours. The entire setup was contained in a sound and light proof behavioural box.

### Behavioural experiments

The experimenter started handling the mice one to two weeks after their arrival. The goal of handling the mice was to familiarize them with the experimenter, experimental room and being moved by hand. We handled mice two or three times, depending on the behaviour of mice, before putting them for the first time in the behavioural arena for one familiarization session.

Each behaviour session started with a shaping period that lasted for ten minutes for the first session and five minutes for the subsequent sessions. In this period, mice could see the grey background on the monitor and explore the arena freely, familiarizing themselves with the environment. After shaping, whenever the mouse was moving, the experimenter started a stimulus. The stimulus passed in the middle along the long axis of the screen in about 3 s, ten times from left-to-right and ten times in the opposite direction, in random order. We showed twenty stimuli in each session with roughly one minute in between. In the session with the habituating and novel stimulus, each stimulus was shown for ten times, in random order. There were one or two days between sessions.

### Stimuli

We started our experiments with a black disc of 1.8 cm in diameter, and a black silhouette of a hawk in flight with a wingspan of 2.4 cm. For investigation of the different features, we also used three other stimuli: 1. black small ellipse that match the hawk’s wingspan and surface area, 2. black small disc matching the surface area of the hawk and small ellipse, 3. black large ellipse with aspect ratio matching the hawk and surface area matching the original black disc. The stimuli were presented used a custom Matlab toolbox built on top of the Psychophysics Toolbox (Kleiner et al., 2007).

### Quantification of freezing

We analysed the video recordings by a custom-made Matlab program, through which we tracked the movements of mice and stimuli, checking for periods of mouse inactivity around the stimulus presentation. We classified as freezing a period of at least 0.5 s during which a mouse decreased its movement to less than 30% of average, starting during the period that the stimulus was visible. A consequence of automatized freezing detection was that some periods of limited movement where mice were not freezing but simply appeared inactive from the vantage point of the camera, were also tagged as freezing. For example, while grooming or sniffing a corner. To overcome these false positives, we monitored each trial and manually corrected the freezing epochs.

### Analysis of head-relative stimulus position

We used a custom Matlab script to determine the position of the snout and the viewing direction of the mouse from the movie. First, the background was automatically determined from movies by taking the mode for each pixel during the whole session. Next, this background was subtracted from each frame and the result was thresholded. After a number of cycles of image dilation and erosion to connect the thresholded regions, the largest connected remaining component was the mouse. The point in this component that was furthest away from its centre-of-mass via an internal path was deemed the tail tip. From the tail tip, the two edges of the tail were traced until they separated more than a certain width. This point was the tail base. The snout was the point on the component that was furthest away from the tail base. The vector from the mouse centre-of-mass to the snout was taken as the approximate head direction. The position from the stimulus was also inferred from the movie. Using these values, we computed the location of the stimulus relative to the mouse head and head-direction.

To compute the conditional probability of a freeze to start in a specific sector (in head-relative coordinates), we computed for each sector in each trial if the stimulus would pass the sector, and if freezing would start while the stimulus was in that sector. These values for each sector were added over all trials of the sessions with only the habituation stimulus for all mice. By dividing these two sums by the total number of sector traversed and freeze starts, we derived the probability that a stimulus traversed a sector, and the probability that a freeze would start in a sector. By dividing the freeze start probability by the traversing probability (adding 0.00001 to the latter, to avoid problems of dividing by zero), we arrived at the conditional probability of a freeze to start in a specific sector.

To compute the probability distribution of freezes when the stimulus was in each sector per mouse (**Fig. 2E**), we counted the number of freezes that occurred in a sector for each mouse and divided by the number of sectors. To compare the observed distribution to a distribution where there is certainly no avoidance of freeze starts in sectors where previous freezes started, we shuffled the trials with freeze starts across the mice, and randomly picked a point on these stimulus trajectories for the freezing to commence.

We also computed the number of freeze starts that occurred in sectors in which the stimulus had previously been the moment a freeze started, or which a stimulus just previously traversed (**Fig. 2F-G)**. This analysis was done across all sessions with one stimulus. To see if this number was lower than the case where mice would have no history of previous stimuli, we again shuffled the trials with freeze starts across the mice, and randomly picked a point on these stimulus trajectories for the freezing to commence.

### Statistical analysis

To see if an individual mouse froze differently between sessions or stimuli, we used a chi^2^ test. To compare freezing averages of a group of mice between sessions or between stimuli, we used paired t-tests. For computing the significance of a possible location-specific effect, we applied 1000 shuffles to the freezing data as described above.

### Data and code availability

All the scripts for stimulus generation and data analysis are available at http://github.com/heimel/InVivoTools. All experimental movies are available by request to the corresponding author.

## Acknowledgements

We thank Mehran Ahmadlou for his help and Christiaan Levelt for sharing equipment and advice. A.T. and J.A.H. were supported by NWO VIDI grant 864.10.010.

## Competing interests

The authors have no competing interests to declare.

## Supplementary figures

**Supplementary Figure 1.**
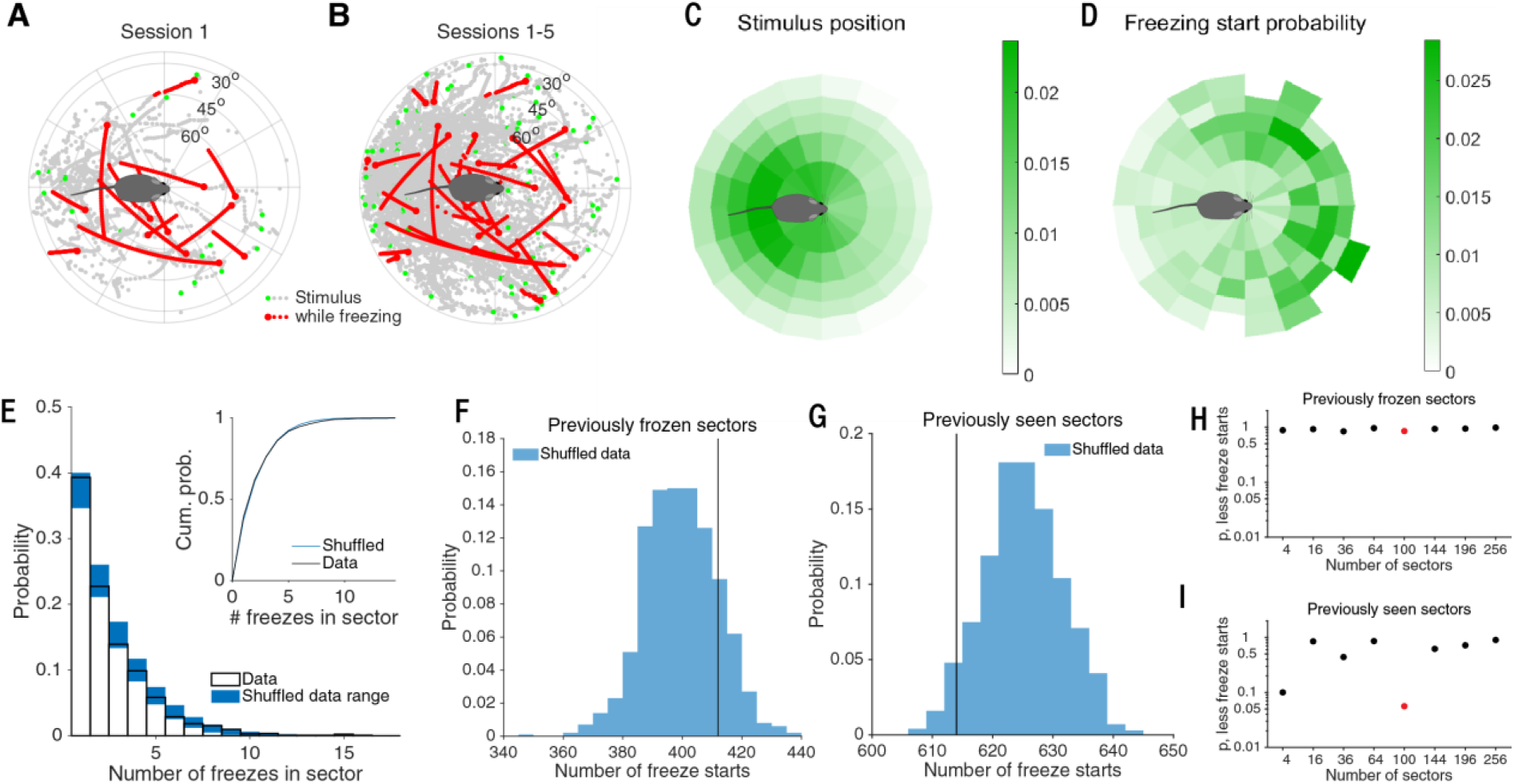
Habituation is not specific to stimulus location. Related to Figure 2. A) Location of the stimulus relative to the mouse for the first session of an example mouse. The entry of the stimulus is shown in green. Red marks the stimulus trajectory while the mouse was freezing. Mouse is not drawn to scale. B) Same as A but for all 5 sessions. C) Probability density of the relative stimulus locations across all mice. Mouse is not drawn to scale. D) The conditional probability of a freeze to start for a given stimulus location. E) Distribution of the number of freezes occurring for each stimulus location for the division in sectors used in C and D. In blue the range of 2 standard deviations below and above the mean for the cases when the freezes are shuffled across mice. Inset shows the cumulative distribution of the data and mean of the shuffled distributions. F) The number of freeze starts that occur in a sector that was previously traversed by the stimulus while a mouse was frozen (black line) falls inside the distribution when the freezes are shuffled across mice. G) Same as G for sectors traversed by the stimulus regardless of whether the mouse was freezing. H) Fraction of number of freeze starts in previously frozen sector of shuffled distribution that is smaller than the observed number. Red marker indicates case shown in F. P-values are not corrected for multiple comparisons. I) Same as H for the number of freeze starts in previously seen sectors. Red marker indicates case shown in G.

## Notes

### Competing Interest Statement

The authors have declared no competing interest.

## References

Bradley, M. M., Lang, P. J., & Cuthbert, B. N. (1993). Emotion, novelty, and the startle reflex: Habituation in humans. Behavioral Neuroscience, 107(6), 970–980. https://doi.org/10.1037/0735-7044.107.6.970

Cooke, S. F., Komorowski, R. W., Kaplan, E. S., Gavornik, J. P., & Bear, M. F. (2015). Visual recognition memory, manifested as long-term habituation, requires synaptic plasticity in V1. Nature Neuroscience, 18(2), 262–271. https://doi.org/10.1038/nn.3920

De Franceschi, G., & Solomon, S. G. (2018). Visual response properties of neurons in the superficial layers of the superior colliculus of awake mouse. Journal of Physiology, 596(24), 6307–6332. https://doi.org/10.1113/JP276964

De Franceschi, G., Vivattanasarn, T., Saleem, A. B., & Solomon, S. G. (2016). Vision Guides Selection of Freeze or Flight Defense Strategies in Mice. Current Biology, 26(16), 2150–2154. https://doi.org/10.1016/j.cub.2016.06.006

Dean, P., Redgrave, P., & Westby, G. W. M. (1989). Event or emergency? Two response systems in the mammalian superior colliculus. Trends in Neurosciences, 12(4), 137–147. https://doi.org/10.1016/0166-2236(89)90052-0

Deecke, V. B., Slater, P. J. B., & Ford, J. K. B. (2002). Selective habituation shapes acoustic predator recognition in harbour seals. Nature, 420, 171–173.

Eilam, D. (2005). Die hard: A blend of freezing and fleeing as a dynamic defense - Implications for the control of defensive behavior. Neuroscience and Biobehavioral Reviews, 29(8), 1181–1191. https://doi.org/10.1016/j.neubiorev.2005.03.027

Evans, D. A., Stempel, A. V., Vale, R., Ruehle, S., Lefler, Y., & Branco, T. (2018). A synaptic threshold mechanism for computing escape decisions. Nature, 558(7711), 590–594. https://doi.org/10.1038/s41586-018-0244-6

Ewert, J. P. (1987). Neuroethology of releasing mechanisms: Prey-catching in toads. Behavioral and Brain Sciences, 10(3), 337–368. https://doi.org/10.1017/S0140525X00023128

Finkenstädt, T., & Ewert, J. P. (1988). Stimulus-specific long-term habituation of visually guided orienting behavior toward prey in toads: a14C-2DG study. Journal of Comparative Physiology A, 163(1), 1–11. https://doi.org/10.1007/BF00611991

Gutfreund, Y. (2012). Stimulus-specific adaptation, habituation and change detection in the gaze control system. Biological Cybernetics, 106(11–12), 657–668. https://doi.org/10.1007/s00422-012-0497-3

Hoy, J. L., Yavorska, I., Wehr, M., & Niell, C. M. (2016). Vision Drives Accurate Approach Behavior during Prey Capture in Laboratory Mice. Current Biology, 26(22), 3046–3052. https://doi.org/10.1016/j.cub.2016.09.009

Kleiner, M., Brainard, D., & Pelli, D. (2007). What’s new in Psychtoolbox-3? Perception, 36.

Lee, K. H., Tran, A., Turan, Z., & Meister, M. (2020). The sifting of visual information in the superior colliculus. ELife, 9. https://doi.org/10.7554/elife.50678

Lissek, S., & van Meurs, B. (2015). Learning models of PTSD: Theoretical accounts and psychobiological evidence. International Journal of Psychophysiology, 98(3), 594–605. https://doi.org/10.1016/j.ijpsycho.2014.11.006

Lorenz, K. (1939). Vergleichende verhaltensforschung. Verhandlungen Der Deutschen Zoologischen Gesellschaft Zoologischer Anzeiger, Supplement(12), 69–102.

McDiarmid, T. A., Bernardos, A. C., & Rankin, C. H. (2017). Habituation is altered in neuropsychiatric disorders—A comprehensive review with recommendations for experimental design and analysis. In Neuroscience and Biobehavioral Reviews (Vol. 80). Elsevier Ltd. https://doi.org/10.1016/j.neubiorev.2017.05.028

Orioli, G., Bremner, A. J., & Farroni, T. (2018). Multisensory perception of looming and receding objects in human newborns. Current Biology, 28(22), R1294–R1295. https://doi.org/10.1016/j.cub.2018.10.004

Peeke, H. V. S., & Veno, A. (1973). Stimulus specificity of habituated aggression in the stickleback (Gasterosteus aculeatus). Behavioral Biology, 8(3), 427–432. https://doi.org/10.1016/S0091-6773(73)80083-5

Prusky, G. T., West, P. W. R., & Douglas, R. M. (2000). Behavioral assessment of visual acuity in mice and rats. Vision Research, 40(16), 2201–2209. https://doi.org/10.1016/S0042-6989(00)00081-X

Randlett, O., Haesemeyer, M., Forkin, G., Shoenhard, H., Schier, A. F., Engert, F., & Granato, M. (2019). Distributed Plasticity Drives Visual Habituation Learning in Larval Zebrafish. Current Biology, 29(8), 1337-1345.e4. https://doi.org/10.1016/j.cub.2019.02.039

Rankin, C. H., Abrams, T., Barry, R. J., Bhatnagar, S., Clayton, D. F., Colombo, J., Coppola, G., Geyer, M. A., Glanzman, D. L., Marsland, S., Mcsweeney, F. K., Wilson, D. A., Wu, C., & Thompson, R. F. (2009). Neurobiology of Learning and Memory Habituation revisited : An updated and revised description of the behavioral characteristics of habituation. Neurobiology of Learning and Memory, 92(2), 135–138. https://doi.org/10.1016/j.nlm.2008.09.012

Salay, L. D., Ishiko, N., & Huberman, A. D. (2018). A midline thalamic circuit determines reactions to visual threat. Nature, 557(7704). https://doi.org/10.1038/s41586-018-0078-2

Schleidt, W., Shalter, M. D., & Moura-Neto, H. (2011). The Hawk/Goose Story: The Classical Ethological Experiments of Lorenz and Tinbergen, Revisited. Journal of Comparative Psychology, 125(2), 121–133. https://doi.org/10.1037/a0022068

Sokolov, E. N. (1963). Higher nervous functions: The orienting reflex. Annual Review of Physiology, 25, 545–580.

Temizer, I., Donovan, J. C., Baier, H., & Semmelhack, J. L. (2015). A Visual Pathway for Looming-Evoked Escape in Larval Zebrafish. Current Biology, 25(14), 1823–1834. https://doi.org/10.1016/j.cub.2015.06.002

Thompson, R. F. (2009). Habituation: A history. Neurobiology of Learning and Memory, 92(2), 127–134. https://doi.org/10.1016/j.nlm.2008.07.011

Thompson, R. F., & Spencer, W. A. (1966). Habituation: A model phenomenon for the study of neuronal substrates of behavior. Psychological Review, 73(1), 16–43. https://doi.org/10.1037/h0022681

Tinbergen, N. (1939). Why do birds behave as they do? (II). In Bird-Lore, 41 (pp. 23–30).

Tovote, P., Esposito, M. S., Botta, P., Chaudun, F., Fadok, J. P., Markovic, M., Wolff, S. B. E., Ramakrishnan, C., Fenno, L., Deisseroth, K., Herry, C., Arber, S., & Lüthi, A. (2016). Midbrain circuits for defensive behaviour. Nature, 534(7606), 206–212. https://doi.org/10.1038/nature17996

Vogel, E. H., & Wagner, A. R. (2005). Stimulus specificity in the habituation of the startle response in the rat. Physiology and Behavior, 86(4), 516–525. https://doi.org/10.1016/j.physbeh.2005.08.042

Yilmaz, M., & Meister, M. (2015). Report Rapid Innate Defensive Responses of Mice to Looming Visual Stimuli. Current Biology, 23(20), 2011–2015. https://doi.org/10.1016/j.cub.2013.08.015

